# Untargeted CUT&Tag and BG4 CUT&Tag are both enriched at G-quadruplexes and accessible chromatin

**DOI:** 10.1101/2024.09.26.615263

**Authors:** Matthew Thompson, Alicia Byrd

**Affiliations:** Department of Biochemistry and Molecular Biology, University of Arkansas for Medical Sciences, Little Rock, AR, 72205, USA; Winthrop P. Rockefeller Cancer Institute, Little Rock, AR, 72205, USA

## Abstract

G-quadruplex DNA structures (G4s) form within single-stranded DNA in nucleosome-free chromatin. As G4s modulate gene expression and genomic stability, genome-wide mapping of G4s has generated strong research interest. Recently, the Cleavage Under Targets and Tagmentation (CUT&Tag) method was performed with the G4-specific BG4 antibody to target Tn5 transposase to G4s. While this method generated a novel high-resolution map of G4s, we unexpectedly observed a strong correlation between the genome-wide signal distribution of BG4 CUT&Tag and accessible chromatin. To examine whether untargeted Tn5 cutting at accessible chromatin contributes to BG4 CUT&Tag signal, we examined the genome-wide distribution of signal from untargeted (i.e. negative control) CUT&Tag datasets. We observed that untargeted CUT&Tag signal distribution was highly similar to both that of accessible chromatin and of BG4 CUT&Tag. We also observed that BG4 CUT&Tag signal increased at mapped G4s, but this increase was accompanied by a concomitant increase in untargeted CUT&Tag at the same loci. Consequently, enrichment of BG4 CUT&Tag over untargeted CUT&Tag was not increased at mapped G4s. These results imply that either the vast majority of accessible chromatin regions contain mappable G4s or that the presence of G4s within accessible chromatin cannot reliably be determined using BG4 CUT&Tag alone.

**GRAPHICAL ABSTRACT:** 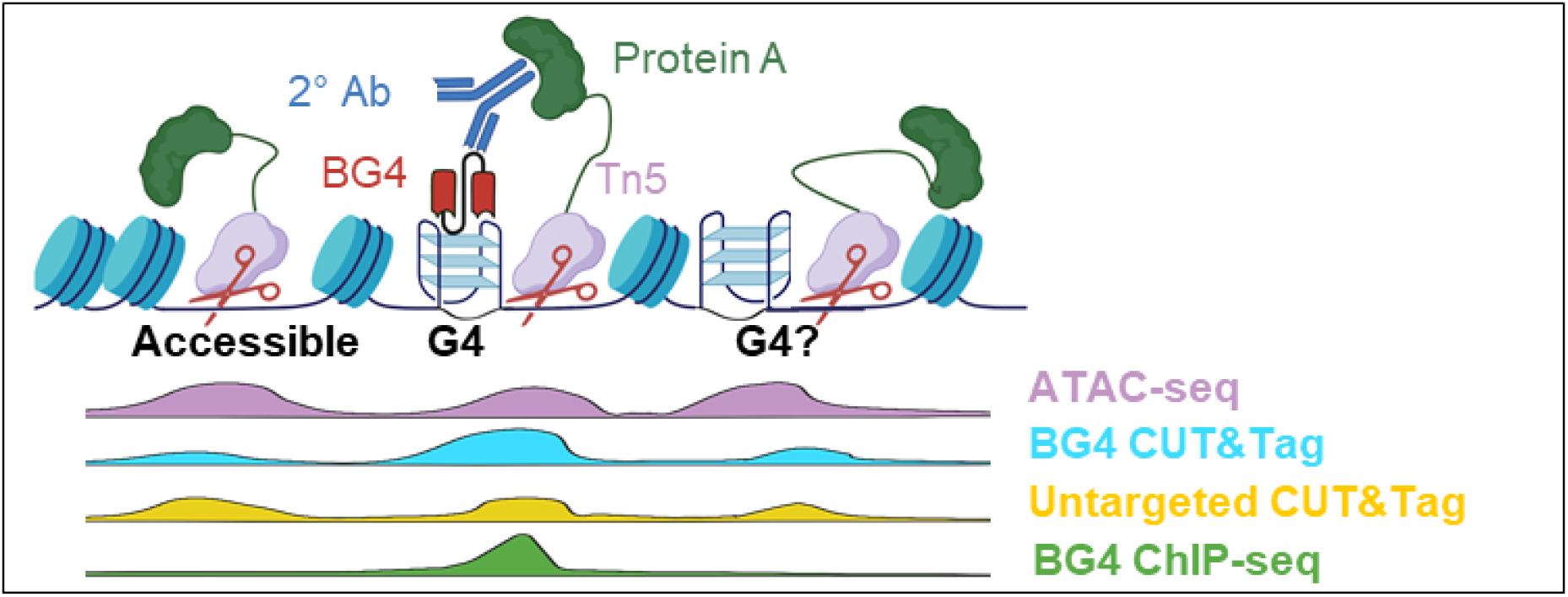

## INTRODUCTION

Formation of non-B DNA secondary structures is both a driver and a consequence of myriad biological processes (reviewed in (1)). G-quadruplex DNA structures (G4s) are present throughout regulatory regions of the human genome (2) and influence a diverse array of cellular processes such as gene expression (3–5), DNA replication progression (6–8), and DNA replication origin firing (9, 10). G4s are formed by runs of multiple guanine bases that interact via Hoogsteen base pairing to form planar tetrads (11) that stack into a G4 structure (12). G4s can subsequently be bound and/or resolved by endogenous proteins to modulate transcription and safeguard genomic stability (reviewed in (13, 14)). Additionally, there exist ongoing efforts to selectively stabilize/destabilize G4s *in cellulo* with chemical ligands to modulate translationally relevant biological processes (reviewed in (15)). Consequently, efforts to identify and map folded G4s throughout the human genome has generated significant research interest (reviewed in (16)).

Initial G4 mapping efforts relied on algorithmic prediction (17–19) and next-generation sequencing methods (20) to identify sequences with the potential to form G4s *in vitro*. However, development of the BG4 single-chain antibody that binds specifically to G4s (21) allowed for genome-wide G4 mapping *in cellulo* using chromatin immunoprecipitation with sequencing (ChIP-seq) (2). Recently, new methods of mapping G4s have been created such as G4Access, which uses controlled micrococcal nuclease (MNase) digestion and size selection to isolate folded G4s within subnucleosomal genomic DNA fragments (22).

Separate efforts to improve the relatively poor signal-to-noise ratio of ChIP-seq for a variety of genomic targets resulted in the creation of another MNase-based method known as Cleavage Under Targets and Release Using Nuclease (CUT&RUN) (23). Further advancement of CUT&RUN then resulted in the creation of the Tn5 transposase-based Cleavage Under Targets and Tagmentation (CUT&Tag) (24, 25) that reduced both the cost and effort of the method when compared to CUT&RUN. CUT&Tag utilizes a fusion protein comprised of the antibody-targeting Protein A (and sometimes Protein G) linked to the Tn5 transposase to selectively fragment and tag (i.e. “tagment”) genomic DNA in the vicinity of an antibody target (24). This antibody-based targeting of Tn5 in CUT&Tag contrasts with the lack of targeting of Tn5 tagmentation in the assay for transposase-accessible chromatin with sequencing (ATAC-seq) that maps nucleosome-depleted accessible chromatin (26).

Recently, CUT&Tag has been adapted to map G4s in bulk and single-cell populations using the BG4 single-chain antibody that binds specifically to G4s (27). The use of BG4 CUT&Tag has identified novel G4s that were not previously observed using BG4 ChIP-seq (27), but this is not wholly unexpected, given the improved signal-to-noise ratio of CUT&Tag compared to ChIP-seq (24). However, use of the Tn5 transposase during CUT&Tag creates a susceptibility to off-target DNA cleavage at open chromatin sites that lack the target of interest (24, 28). This could result in CUT&Tag signal accumulation at Tn5-preferred sites under untargeted conditions (i.e. using non-targeting IgG or omitting IgG).

Recent literature has highlighted the utility of identifying regions of the genome to which an abundance of untargeted sequencing reads map. The genome-wide distributions of untargeted signal enrichment in ChIP-seq data (29) and in CUT&RUN data (30) have been generated, and these distributions reflect genomic assembly artefacts and highlight biases intrinsic to the differing methodologies (i.e. DNA shearing via sonication in ChIP-seq and digestion with MNase in CUT&RUN). To our knowledge, no similar map of problematic regions of the genome related to CUT&Tag analyses has been generated.

Given that G4s require an open chromatin environment to form, we hypothesized that Tn5-preferred cleavage at open chromatin during CUT&Tag could generate local CUT&Tag signal enrichment in the vicinity of G4s. To test this, we compiled untargeted CUT&Tag datasets and assessed the local sequence and chromatin environment at sites of signal enrichment in untargeted CUT&Tag. We additionally compared enrichment of BG4 CUT&Tag over untargeted CUT&Tag at mapped G4s in multiple cell lines to assess the capability of BG4 CUT&Tag to specify the presence or absence of G4s within open chromatin, despite the presence of untargeted CUT&Tag signal enrichment at these sites. We observed that although the overall amplitude of BG4 CUT&Tag signal increases at mapped G4s, the enrichment of BG4 CUT&Tag above untargeted CUT&Tag is not increased in the presence of a mapped G4 within regions of accessible chromatin. These results suggest that reliance solely on peaks within BG4 CUT&Tag data to map G4s can lead to an increase in false negative and/or false positive identification of G4s throughout the genome.

## MATERIAL AND METHODS

### Read mapping and peak calling

Untargeted CUT&Tag datasets in the Gene Expression Omnibus (GEO) (31) were queried using the following: ((CUT AND Tag AND “homo sapiens”)) AND “.bw” AND ((“gse”[Filter] OR “gds”[Filter] OR “gsm”[Filter]) AND “Homo sapiens”[Organism]). Untargeted CUT&Tag reads (**Supplementary Table 1**) and reads from other data sources (**Supplementary Tables 2 and 3**) were downloaded from the Sequence Read Archive (SRA) (11) as .fastq files using fastq-dump (https://github.com/ncbi/sra-tools) or were downloaded from ENCODE (29) as .bam files pre-mapped to the GRCh38 assembly of the human genome (33). Raw reads (.fastq) were filtered (fastqc), trimmed (34), and aligned (35) to the GRCh38 assembly of the human genome using the Nextflow (36, 37) cutandrun pipeline, v3.2.1 (38). Read counts were normalized by read depth with counts-per-million reads (CPM) normalization with a bin size of 50, and peaks were called with MACS2 (39) (default settings) or with SEACR (40) with a threshold of 0.02. For ENCODE-sourced reads (**Supplementary Table 3**) already mapped to GRCh38, reads were indexed, sorted, and normalized by CPM using deeptools (41) bamCoverage with an effective genome size of 2913022398 and a bin size of 50. When available, peaks of signal enrichment that were called and previously analysed in the source publication were utilized. Unless otherwise indicated, ATAC-seq, CUT&Tag, ChIP-seq, and G4Access datasets were generated in K562 cells (**Supplementary Tables 2 and 3**).

### Peak overlap analysis

Genomic intervals present in peaks between more than two datasets were identified using bedtools (42) multiinter, whereas shared or unshared peaks from two datasets were identified using GenomicRanges (43). Read counts at individual peaks were quantified using the UCSC Genome Browser (44) utility bigWigAverageOverBed by quantifying the average normalized read count signal over each base within a peak with non-covered bases being quantified with a signal of zero.

### Peak annotation and motif finding

Peaks in untargeted CUT&Tag datasets were annotated using ChIPseeker (45) annotatePeak or with HOMER (46) annotatePeaks with default settings. Motifs were found with HOMER findMotifsGenome with a motif size of 20.

### Peak visualization

Heatmaps and metaplots of normalized read counts centred at peaks were generated using deeptools (41) computeMatrix and plotHeatmap. Normalized read counts were visualized across the genome using IGV (47).

## RESULTS

### Untargeted CUT&Tag samples have a characteristic pattern of genome-wide enrichment

We acquired untargeted CUT&Tag datasets from publicly available data (**Table 1**) to assess whether these data display any pattern of consistent genome-wide enrichment. We selected datasets originating from a variety of cell lines and labs to capture dataset- and cell line-agnostic signal enrichment. Raw reads were reprocessed using nextflow cutandrun (38), and peaks were called using both MACS2 (39) (default settings) and SEACR (40) (top 2% of peaks). The high threshold for SEACR was utilized to capture a similar number of peaks as with MACS2 in order to capture genomic regions with read counts which were globally high (SEACR) and/or locally distinguishable (MACS2). We defined consistent signal enrichment as any called peak from either peak caller which was present in more than 33.3% of datasets (at least 7 of 21; similar to ref (30)). 5,163 regions of consistent signal enrichment were identified (**Figure 1A**). This set of conserved peaks is subsequently referred to as untargeted CUT&Tag peaks. The majority of peaks called are shared between both peak calling algorithms (**Supplementary Figure 1**), indicating that untargeted CUT&Tag peaks are frequently both high in amplitude and enriched above local background.

**Figure 1.**
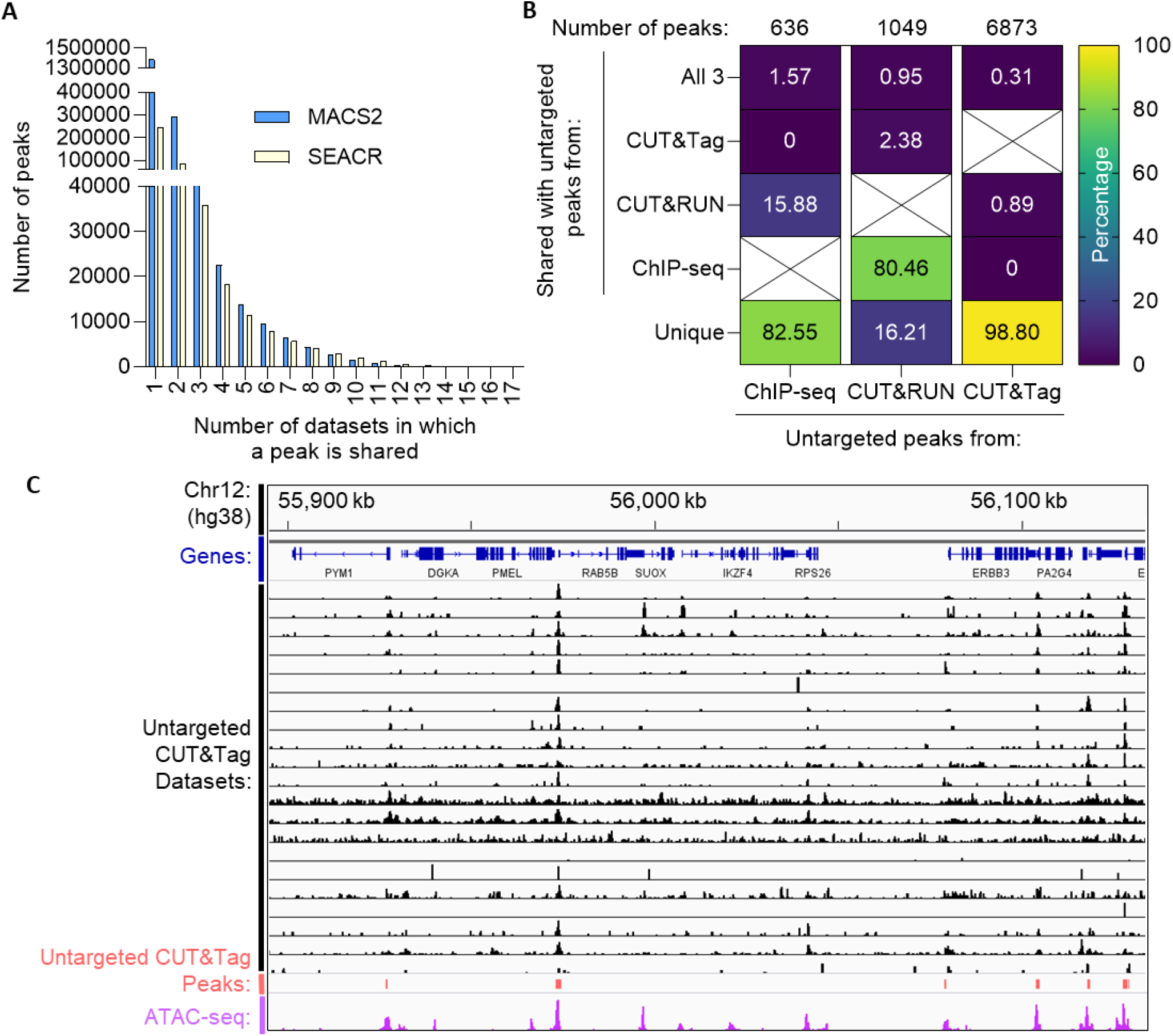
Untargeted CUT&Tag has a consistent genome-wide enrichment. (**A**) Histogram of the number of peaks called by either MACS2 or SEACR that are shared by the specified number of datasets. (**B**) Percentage of unique or shared peaks between peaks of enrichment in untargeted ChIP-seq (48), untargeted CUT&RUN (30), and untargeted CUT&Tag samples. (**C**) Read counts for untargeted CUT&Tag datasets at a representative locus on chromosome 12 overlaid onto the untargeted CUT&Tag consensus peaks and ATAC-seq signal (average of 3 biological replicates) from K562 cells.

Untargeted CUT&Tag peaks have a unique genome-wide distribution compared to regions of signal enrichment from untargeted ChIP-seq data (48) and untargeted CUT&RUN data (30) (**Figure 1B**). The high proportion of genomic regions of untargeted CUT&RUN signal enrichment that overlap untargeted ChIP-seq enrichment (80.46%) has been determined to originate from artefactual mapping of short reads to the GRCh38 assembly, where the remaining unshared peaks of untargeted CUT&RUN enrichment (16.21%) were hypothesized to originate from biases in MNase-catalyzed DNA cleavage (30). Similarly, the abundance of unique peaks identified in untargeted ChIP-seq data that do not overlap untargeted CUT&RUN or CUT&Tag peaks (82.55%) are hypothesized to reflect biases related to crosslinking and sonication. Following the same logic, the high percentage of unique peaks (98.80%) in untargeted CUT&Tag data and the low percentage of peaks from untargeted CUT&RUN (1.57%) and ChIP-seq (3.33%) datasets that overlap untargeted CUT&Tag peaks suggest that the observed signal enrichment may be due to DNA sequence or DNA structural biases in Tn5-catalyzed cleavage. This suggested to us that the genome-wide distribution of untargeted CUT&Tag enrichment could resemble the Tn5-based ATAC-seq method which maps accessible chromatin (26). Indeed, untargeted CUT&Tag peaks frequently overlapped sites of ATAC-seq enrichment in K562 cells (**Figure 1C**), suggesting an enrichment at accessible chromatin and regulatory regions.

### Untargeted CUT&Tag peaks are enriched at Tn5-accessible regulatory regions

To quantify the overlap of untargeted CUT&Tag peaks with ATAC-seq-mapped regulatory regions, we first annotated the set of untargeted CUT&Tag consensus peaks. A high percentage (93.94%) of untargeted CUT&Tag peaks are located within 3 kb of a promoter, with 91.13% of peaks being located less than 1 kb from a promoter (**Figure 2A**). A separate annotation corroborated this, as 79.68% of peaks are annotated as being at a regulatory region, either at a promoter (68.41%) or CpG island (11.27%; **Figure 2B**). Additionally, quantification of signal from ATAC-seq from a representative ENCODE Consortium dataset revealed that ATAC-seq signal increases within 3 kb of a majority of untargeted CUT&Tag peaks (**Figure 2C**). A similar pattern of enrichment was not observed within 3 kb of untargeted ChIP-seq or CUT&RUN peaks (**Figure 2D-E**). This demonstrates that untargeted CUT&Tag peaks colocalize with Tn5-accessible chromatin within regulatory regions.

**Figure 2.**
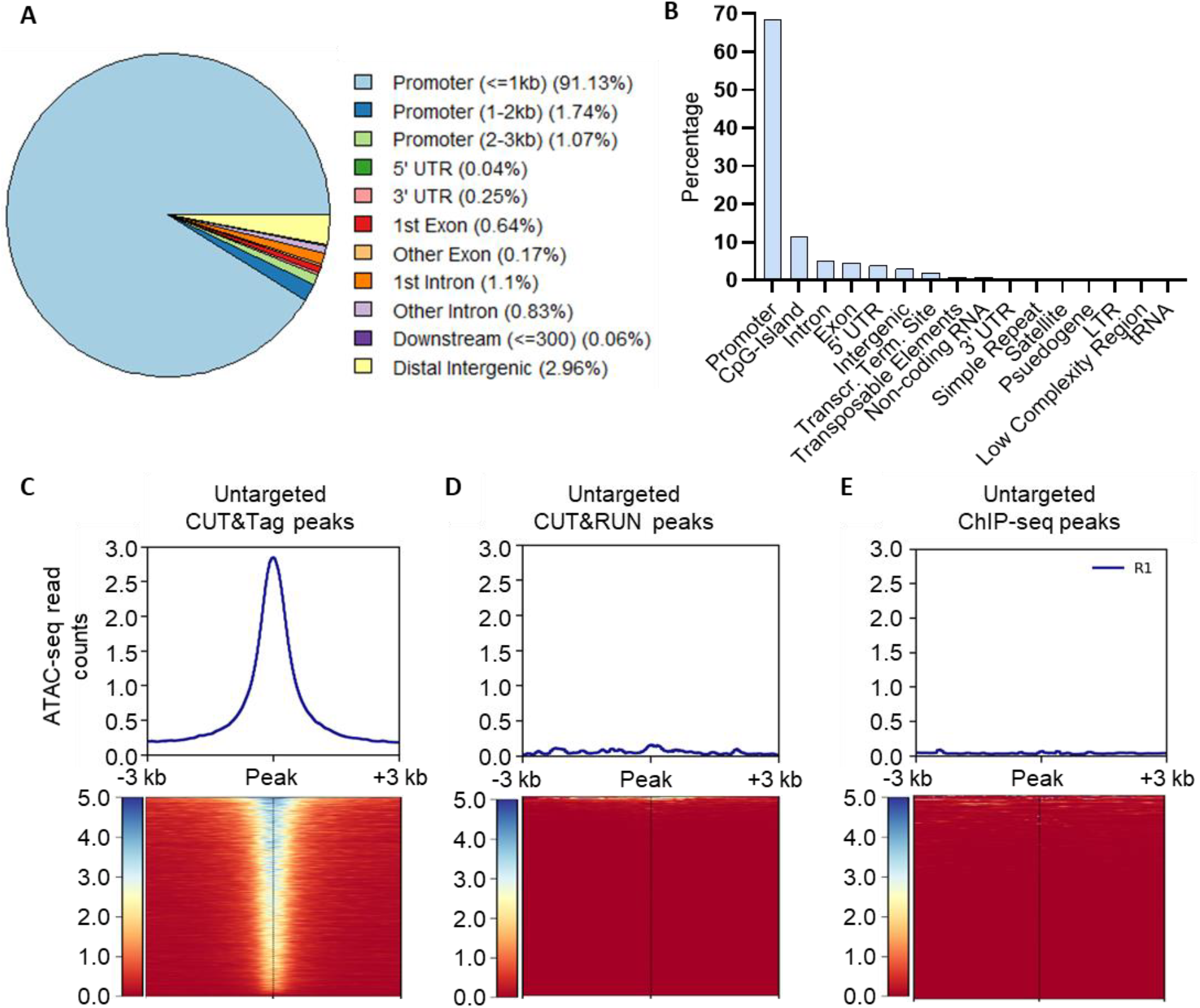
Untargeted CUT&Tag peaks colocalize with regulatory regions and accessible chromatin. (**A**) Proportions of annotations overlapping with untargeted CUT&Tag peaks using ChIPseeker (45). (**B**) Proportions of annotations overlapping with untargeted CUT&Tag peaks using HOMER (46). Metaplots of ATAC-seq signal (ENCSR868FGK; average of 3 biological replicates) at peaks of signal enrichment in untargeted CUT&Tag (**C**), CUT&RUN (30) (**D**), and ChIP-seq (48) (**E**) samples.

### Untargeted CUT&Tag peaks colocalize with G4s

Given that accessible regulatory regions of the human genome have the propensity to form DNA secondary structures such as G4s (2), we wondered whether CUT&Tag problematic regions colocalized with G4s. Identification of known sequence motifs that occur in untargeted CUT&Tag peaks revealed the prevalence of motifs with G- and C-rich sequences with tandem runs of guanines or cytosines (**Figure 3A**). Additionally, the untargeted CUT&Tag peaks contain known binding motifs for the Sp1 transcription factor, a *bone fide* G4-binding protein (3, 4). *De novo* motifs also include Sp1 motifs, motifs for the YY1 G4-binding transcription factor (49), and tandem runs of guanines and cytosines (**Figure 3B**). *De novo* motifs also contain TTGxxG/C motifs, which have been previously identified at Tn5 insertion sites (50). The proportion of untargeted CUT&Tag peaks that overlap a G4 was calculated for each set of untargeted CUT&Tag peaks that are shared by 1 or more datasets. The least reproducible untargeted CUT&Tag peaks (present in 4 or fewer datasets) have a low proportion of peaks that overlap a mapped G4 (**Figure 3C**). However, as peak reproducibility increases, the proportion of untargeted CUT&Tag peaks that overlap a mapped G4 also increases, reaching saturation when peaks are shared by at least 5 datasets (**Figure 3C**). This indicates that untargeted CUT&Tag signal is reproducibly enriched at G4s that are algorithmically predicted (pqsfinder) (51), able to form *in vitro* (G4-seq) (52), and observed in human cell lines through use of G4Access (22), BG4 CUT&Tag (27), and/or BG4 ChIP-seq (53).

**Figure 3.**
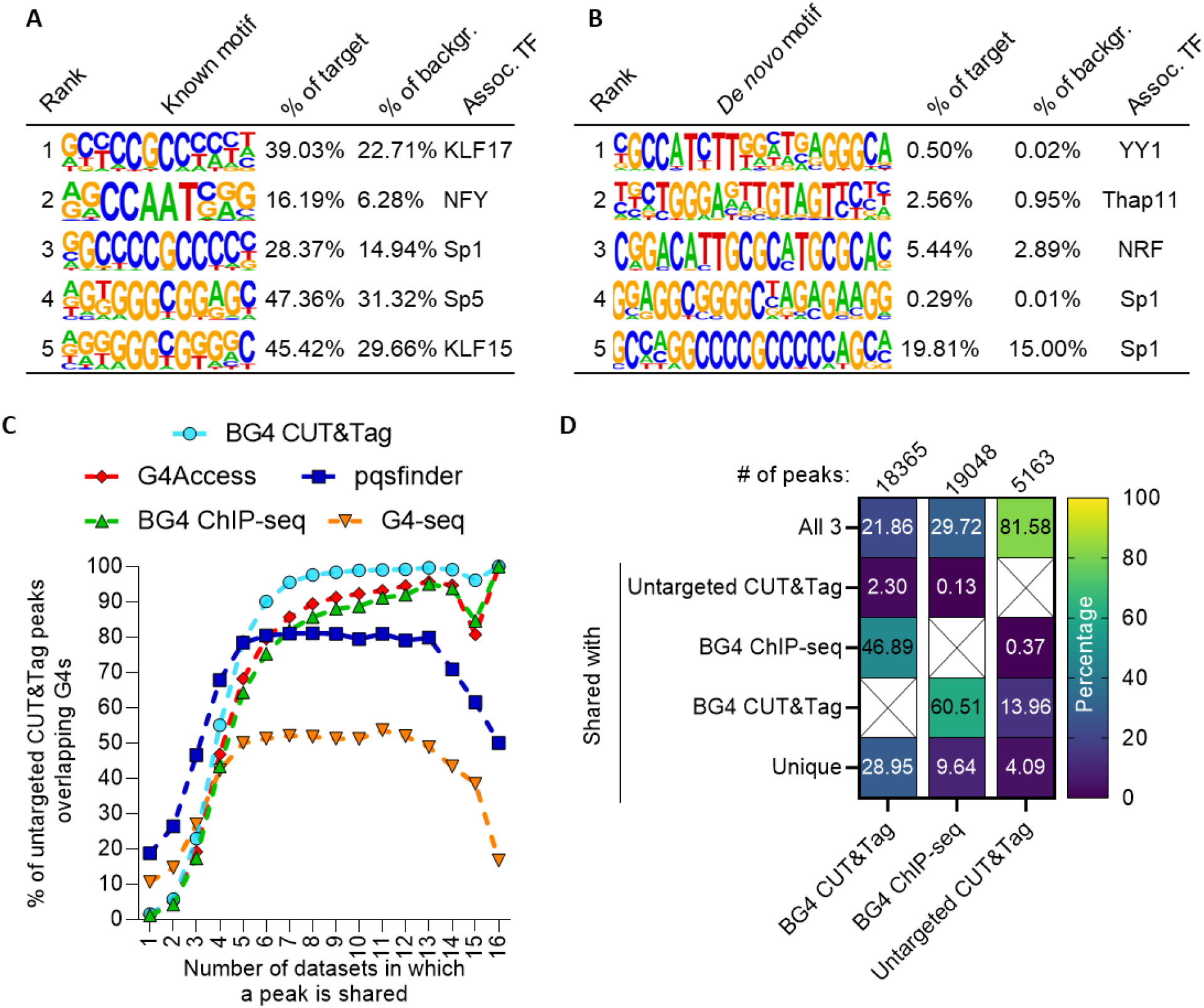
Untargeted CUT&Tag peaks colocalize with mapped G4s. (**A**) Top 5 known motifs and associated transcription factors (TF) present in untargeted CUT&Tag peaks identified by HOMER (46). All p-values < 1×10^-125^ by binomial test. (**B**) Top 5 *de novo* motifs and associated transcription factors (TF) present in untargeted CUT&Tag peaks identified by HOMER (46). All p-values < 1×10^-20^ by binomial test. (**C**) Proportion of untargeted CUT&Tag peaks present in the specified number of datasets that overlap a G4 mapped by BG4 CUT&Tag (27), BG4 ChIP-seq (53), G4Access (22), or mapped *in vitro* by G4-seq (52) or *in silico* by pqsfinder (51). (**D**) Proportion of overlap between untargeted CUT&Tag peaks and BG4 CUT&Tag (27) and/or BG4 ChIP-seq (53).

Additionally, we quantified the degree of colocalization between the set of untargeted CUT&Tag peaks (present in at least 7 of 21 datasets) with two different BG4-based methods of mapping G4s. The majority (95.91%) of the untargeted CUT&Tag peaks overlap G4s mapped by BG4 ChIP-seq (53) and/or BG4 CUT&Tag (27) (**Figure 3D**). However, 13.96% of untargeted CUT&Tag peaks colocalize with only BG4 CUT&Tag G4s and not with BG4 ChIP-seq G4s.

### Enrichment of BG4 CUT&Tag over untargeted CUT&Tag is not increased at G4s

Given our prior observation of colocalized enrichment of both BG4 CUT&Tag and untargeted CUT&Tag, we hypothesized that untargeted Tn5 cutting at accessible chromatin could contribute to signal enrichment in CUT&Tag regardless of the inclusion of the BG4 antibody. This could result in a confounded measurement of the presence of a folded G4 *in cellulo* when using BG4 CUT&Tag, as untargeted CUT&Tag signal could also be enriched at accessible chromatin in the presence of a mapped G4.

To test this hypothesis, , we compared the enrichment of BG4 CUT&Tag signal over untargeted CUT&Tag at mapped G4s (22, 27, 51–53). We reasoned that, if BG4 CUT&Tag signal is derived from BG4 affinity to G4s and is not derived from untargeted cutting at accessible chromatin, then we should observe an increased enrichment of BG4 CUT&Tag above untargeted CUT&Tag signal at G4s that is not observed in the absence of a G4.

Therefore, we first compared the distribution of read counts of BG4 CUT&Tag and untargeted CUT&Tag signal at BG4 CUT&Tag peaks (27) that overlap or do not overlap a mapped G4. Both BG4 CUT&Tag and untargeted CUT&Tag read counts (27) increase at mapped G4s, with the strongest increase at G4s that were mapped *in cellulo* by BG4 ChIP-seq (53) (**Figure 4A, Supplementary Figure 2A**) and G4Access (22) (**Figure 4B**). Additionally, both BG4 CUT&Tag and untargeted CUT&Tag read counts increase at G4s mapped *in vitro* with G4-seq (52) (**Figure 4C, Supplementary Figure 2B**) and *in silico* with pqsfinder (51) (**Figure 4D, Supplementary Figure 2C**).

**Figure 4.**
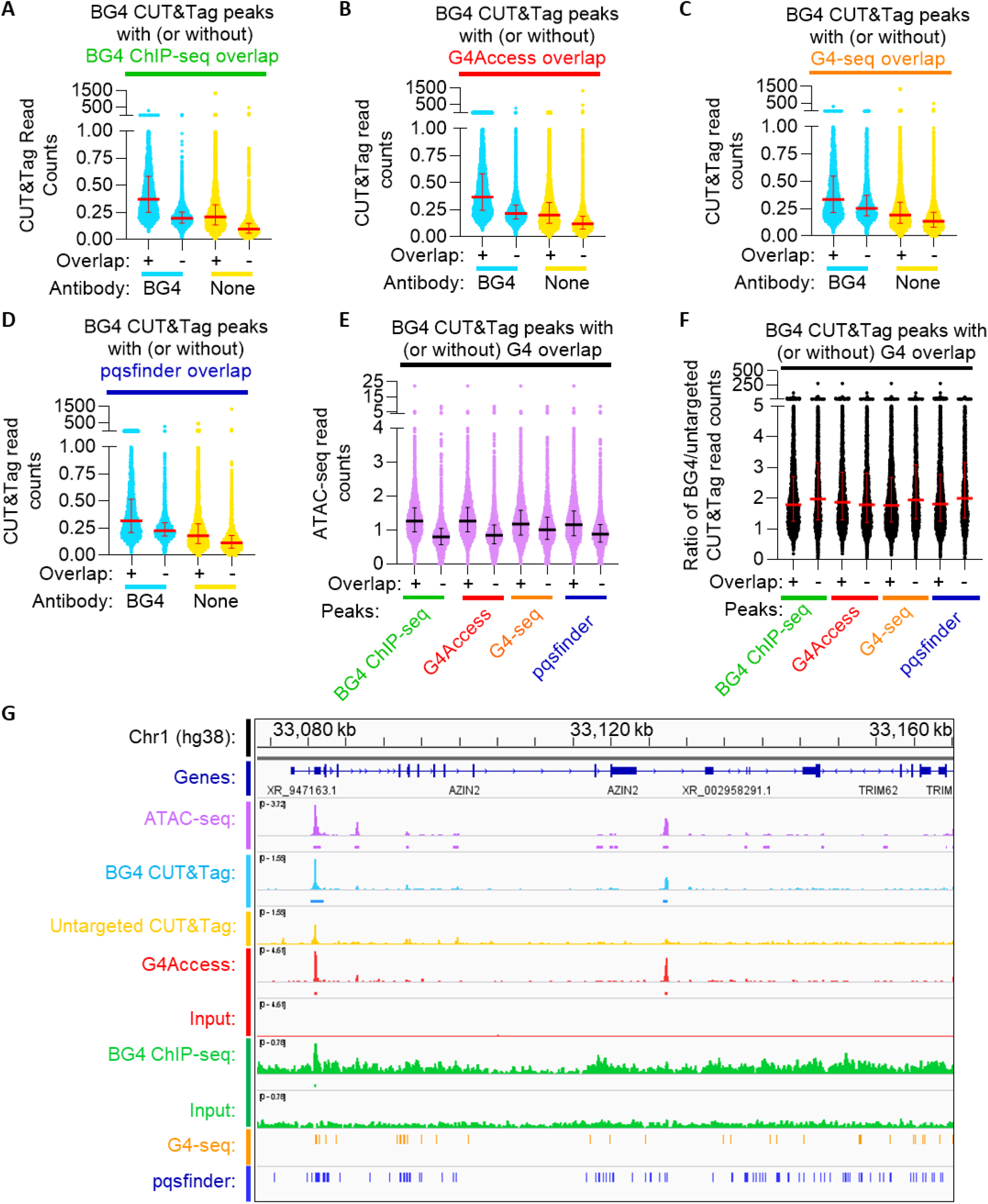
Enrichment of BG4 CUT&Tag over untargeted CUT&Tag does not increase at G4s in K562 cells. (**A-D**) BG4 or untargeted CUT&Tag signal (27) read counts were quantified at BG4 CUT&Tag peaks that overlap or do not overlap a G4 mapped by BG4 ChIP-seq (53) (**A**), G4Access (22) (**B**), G4-seq (52) (**C**), or pqsfinder (51) (**D**) in K562 cells. Data are the average of 3 biological replicates, and the median and interquartile ranges are plotted in red. (**E**) ATAC-seq read counts at BG4 CUT&Tag peaks that overlap or do not overlap a G4 mapped by the indicated method are plotted. Data are the average of 3 biological replicates, and the median and interquartile ranges are plotted in black. (**F**) Ratio of read counts of BG4 CUT&Tag (27) and untargeted CUT&Tag (27) quantified at individual CUT&Tag peaks that overlap or do not overlap a G4 mapped by the indicated method. Data are the average of 3 biological replicates, and the median and interquartile ranges are plotted in red. (**G**) Read counts (signal traces) and called peaks (rectangles) of ATAC-seq, BG4 CUT&Tag (27), untargeted CUT&Tag (27), G4Access (22), G4Access input (22), BG4 ChIP-seq (53) and ChIP-seq input (53) from K562 cells with G4-seq (52) and pqsfinder-predicted G4s (51) at a representative locus on chromosome 1. ATAC-seq, BG4 CUT&Tag, and untargeted CUT&Tag reads are the average of 3 biological replicates, G4Access reads are the average of 2 biological replicates, and G4Access input, BG4 ChIP-seq, and ChIP-seq input are data from 1 biological replicate. BG4 ChIP-seq peaks are the average of two biological replicates.

The concomitant increase in signal for both BG4-targeted and untargeted CUT&Tag suggests a possible increase in chromatin accessibility at these sites, as Tn5 cutting at accessible chromatin can occur in targeted or untargeted CUT&Tag reactions, although the signal amplitude due to Tn5 cutting at off-target accessible chromatin is generally low (24). In line with this, an increase in ATAC-seq signal is observed at the previously examined BG4 CUT&Tag peaks that overlap a mapped G4 (**Figure 4E, Supplementary Figure 2D**). Given that G4 formation requires nucleosome-depleted ssDNA, this is expected. However, it is important to ensure that the increase in BG4 CUT&Tag signal at G4s is related to BG4 antibody-based targeting of Tn5 to G4s (21) and does not occur due the local increase in chromatin accessibility.

If BG4 CUT&Tag signal reflects BG4 affinity and not chromatin accessibility, we would expect to observe an increase in BG4 CUT&Tag signal greater in magnitude than that of untargeted CUT&Tag (i.e. increased enrichment over background) at mapped G4s. To measure this, we quantified the ratio of BG4 CUT&Tag signal over untargeted CUT&Tag signal at each of the individual CUT&Tag peaks that overlap or do not overlap a mapped G4. This reveals a median two-fold enrichment of BG4 CUT&Tag read counts above untargeted CUT&Tag read counts at BG4 CUT&Tag peaks that do not overlap a mapped G4. Unexpectedly, this two-fold ratio of BG4-targeted CUT&Tag over untargeted CUT&Tag does not appreciably change in the presence of a mapped G4 (**Figure 4F, Supplementary Figure 2E**). In fact, for BG4 ChIP-seq (53), G4-seq (52), and pqsfinder (51), the median ratio of BG4 CUT&Tag signal over untargeted CUT&Tag signal at individual CUT&Tag peaks is greater at regions that do not overlap a mapped G4 than those that do overlap a mapped G4. This suggests that CUT&Tag reads from untargeted Tn5 cutting at accessible chromatin strongly contribute to BG4 CUT&Tag signal in addition to the contribution of CUT&Tag reads from BG4-targeted Tn5 cutting at folded G4s *in cellulo*. Supporting this suggestion, CUT&Tag signal enrichment for both BG4 CUT&Tag and untargeted CUT&Tag can be observed at ATAC-seq sites in the absence of a visible enrichment in BG4 ChIP-seq signal (**Figure 4G, Supplementary Figure 2F**). G4Access signal is also observed at these sites, but we also observed a subset of these sites which lack overlap with either a G4-seq G4 (**Figure 4G**) or both G4-seq and pqsfinder G4s (**Supplementary Figure 2F**). Two possible explanations for these results are: BG4 CUT&Tag and G4Access have increased resolution over BG4 ChIP-seq, giving them signal enrichment at G4s that BG4 ChIP-seq is unable to detect, or CUT&Tag and G4Access both share a similar background resulting from enzymatic (Tn5 or MNase) cutting at accessible chromatin. To test these possibilities, we performed an analysis of CUT&Tag signal enrichment over background at accessible chromatin sites in the presence or absence of a mapped G4.

### Enrichment of BG4 CUT&Tag over untargeted CUT&Tag is not increased at G4s within accessible chromatin

To investigate whether CUT&Tag data is enriched at antibody targets compared to all accessible chromatin, we examined enrichment of target CUT&Tag data over untargeted CUT&Tag data at ATAC-seq peaks that overlap or do not overlap ChIP-seq peaks of the CUT&Tag target. Using histone post-translational modifications (PTMs) as controls, we observed a dramatic increase in CUT&Tag read counts (24) at ATAC-seq peaks that overlap the H3K4me3 or H3K27me3 ChIP-seq peaks compared to ATAC-seq peaks that do not overlap a ChIP-seq peak (**Supplementary Figure 3A**). The CUT&Tag read counts at ATAC-seq peaks that do not overlap the appropriate histone PTM ChIP-seq peaks are close to zero for both H3K4me3 and H3K27me3 (**Supplementary Figure 3A**). Additionally, enrichment of target CUT&Tag signal compared to untargeted CUT&Tag is observed at ATAC-seq peaks that overlap the appropriate ChIP-seq peak, as opposed to a lack of enrichment at ATAC-seq peaks which do not overlap a ChIP-seq peak (**Supplementary Figure 3B**). These data demonstrate that for histone PTMs, targeted CUT&Tag signal is significantly enriched above untargeted CUT&Tag at the expected ChIP-seq mapped sites and likely reflects antibody-targeted Tn5 cleavage.

To investigate whether BG4 CUT&Tag signal is enriched over untargeted CUT&Tag at G4s specifically and not at all accessible chromatin, the same analysis was performed. Read counts of BG4 CUT&Tag (27) and untargeted CUT&Tag (27) were examined at ATAC-seq sites that overlap a mapped G4 (22, 51–53). Both BG4 CUT&Tag and untargeted CUT&Tag reads increase at ATAC-seq peaks that overlap a mapped G4 (**Figure 5A, Supplementary Figure 4A**). However, the enrichment of BG4 CUT&Tag over untargeted CUT&Tag does not appreciably increase at mapped G4s (**Figure 5B, Supplementary Figure 4B**). These results suggest that enrichment of BG4 CUT&Tag signal over untargeted CUT&Tag does not necessarily signal the presence of a G4 within accessible chromatin. Indeed, we again observed enrichment of BG4 CUT&Tag (27), untargeted CUT&Tag (27), and G4Access (22) at accessible chromatin in the absence of an accompanying BG4 ChIP-seq mapped G4, but this enrichment was not observed for CUT&Tag reactions targeting histone PTMs (**Figure 5C**). In total, these results suggest that the genome-wide distribution of reads generated by BG4 CUT&Tag and untargeted CUT&Tag are strongly influenced by susceptibility of the local chromatin environment to untargeted Tn5 cutting.

**Figure 5.**
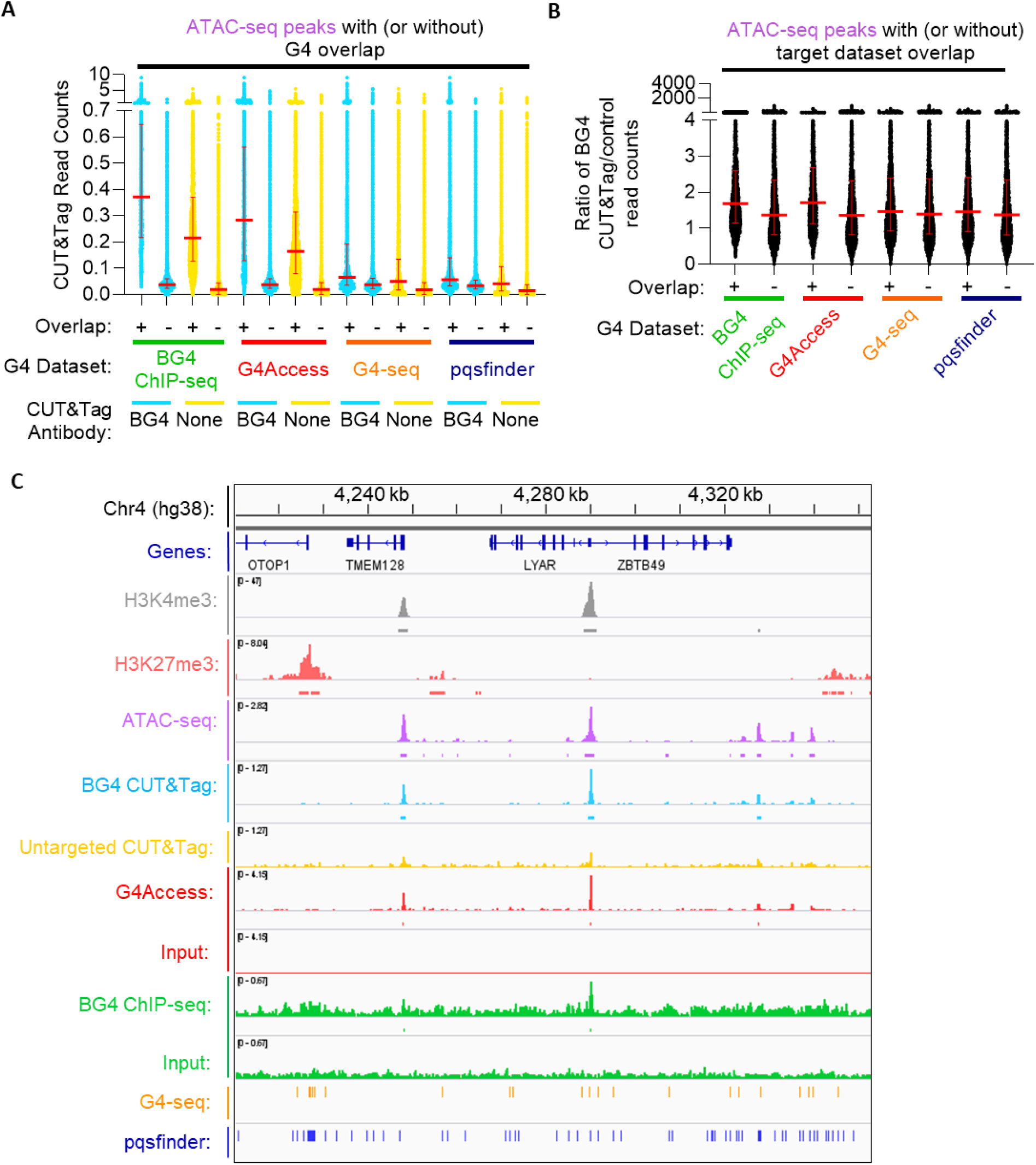
Enrichment of BG4 CUT&Tag over untargeted CUT&Tag does not increase at G4s within accessible chromatin in K562 cells. (**A**) Read counts of BG4 CUT&Tag (27) and untargeted CUT&Tag (27) from K562 cells at ATAC-seq peaks that overlap or do not overlap a mapped G4 (22, 51–53). Data are the average of 3 biological replicates, and the median and interquartile ranges are plotted in red. (**B**) Ratio of read counts of BG4 CUT&Tag (27) and untargeted CUT&Tag (27) from K562 cells quantified at ATAC-seq peaks that overlap or do not overlap a G4 mapped by the indicated method (22, 51–53). Data are the average of 3 biological replicates, and the median and interquartile ranges are plotted in red. (**C**) Read counts (signal traces) and called peaks (rectangles) of H3K4me3 CUT&Tag (24), H3K27me3 CUT&Tag (24), ATAC-seq, BG4 CUT&Tag (27), untargeted CUT&Tag (27), G4Access (22), G4Access input (22), BG4 ChIP-seq (53), and ChIP-seq input (53) from K562 cells with G4-seq (52) and pqsfinder-predicted G4s (51) at a representative locus on chromosome 4. Read counts of H3K4me3 CUT&Tag, H3K27me3 CUT&Tag, G4Access input, BG4 ChIP-seq, and ChIP-seq input are from 1 biological replicate. Read counts from G4Access are the average of 2 biological replicates. Read counts from ATAC-seq, BG4 CUT&Tag, and untargeted CUT&Tag are the average of 3 biological replicates. BG4 ChIP-seq peaks are the average of two biological replicates.

## DISCUSSION

The need to predict and map G4s (reviewed in (16)) has spurred technical innovation, and a diverse suite of G4 mapping tools exists to meet multiple experimental needs. For example, full-length antibodies (54), single-chain antibodies (21), nanobodies (55), protein-based probes (56), and G4-ligand-based probes (57) have been utilized for chromatin immunoprecipitation-(58) and *in situ* Tn5-based sequence mapping methods (27, 57). Additionally, the G4Access method uses the G-rich sequence cutting preference of MNase combined with the inhibition of MNase cutting by nucleosomes and folded G4s to map G4s within accessible chromatin (22). While each method has limitations, BG4 CUT&Tag (27) has been noted for its high resolution compared to BG4 ChIP-seq and low input requirements (24) which can enable G4 mapping of single cell populations (27) and precious patient-derived samples. However, CUT&Tag analyses can be influenced by untargeted Tn5 cutting at accessible chromatin (24, 28). Indeed, simply by reducing the ionic strength of the buffer during tagmentation, Tn5-based cleavage can be used to map accessible chromatin in the vicinity of antibody targets, rather than solely mapping the site of antibody binding (59). Given the dynamic nature of G4s and their high degree of overlap with accessible regulatory chromatin, it is important to characterize which G4 mapping methods can be used to distinguish the presence or absence of G4s without mapping accessible chromatin environments instead.

The initially-reported method to distinguish between antibody-targeted and untargeted Tn5 tagmentation at accessible chromatin for CUT&Tag analyses used read count-based discrimination to separate high read count targeted signals from low read count untargeted signals (24). The read count difference between targeted CUT&Tag of histone PTMs compared to untargeted Tn5 tagmentation is sufficient to discriminate between on- and off-target CUT&Tag signal enrichment (**Supplementary Figure 3A**). For peaks of BG4 CUT&Tag signal with the highest read count enrichment, this method could be employed to identify high-confidence peaks of CUT&Tag enrichment. Indeed, this method has been utilized to call regions with the top 5% of signal using SEACR (40) without a negative control dataset (27). However, the high degree of overlap between the read count distributions of BG4-targeted and untargeted CUT&Tag combined with the simultaneous enrichment of both BG4-targeted and untargeted CUT&Tag signals at mapped G4s (**Figure 4**) would create a large population of false negative or false positive G4s using this method of read count discrimination. Additionally, the simultaneous enrichment of both BG4-targeted and untargeted CUT&Tag reads at mapped G4s and accessible chromatin may increase false negative results (i.e. lack of a peak call) when a negative control is included during peak calling, as an increase in targeted BG4 CUT&Tag signal at a folded G4 could be masked by the simultaneous increased read counts of untargeted CUT&Tag at the same locus.

Consequently, the results of our analysis place limitations on identifying targeted signal enrichment at antibody targets over untargeted enrichment at accessible chromatin using CUT&Tag. This limitation poses minimal issues for CUT&Tag of highly abundant, high-occupancy targets such as histone PTMs, as enrichment of target CUT&Tag signal over untargeted CUT&Tag signal strongly increases at mapped target sites within accessible chromatin (**Supplementary Figure 3**). However, BG4-targeted CUT&Tag for G4s does not result in an appreciable increase in signal enrichment over untargeted signal enrichment at mapped G4s within accessible chromatin (**Figure 5 and Supplementary Figure 4**).Additionally, we observed the presence of BG4 CUT&Tag signal enrichment at accessible chromatin that is not readily observed using BG4 ChIP-seq (53) (**Figure 5C**).

The lack of a strong increase in BG4 CUT&Tag enrichment over untargeted CUT&Tag at mapped G4s and the identification of novel BG4 peaks of enrichment when compared to BG4 ChIP-seq (53) could be the result of a combination of multiple factors. First, Tn5-based DNA cleavage methods result in an increase in signal-to-noise when compared to BG4 ChIP-seq, so BG4 CUT&Tag signal enrichment at sites lacking visible BG4 ChIP-seq enrichment is not unexpected. Second, Tn5-based methods can result in untargeted cleavage at accessible chromatin (24, 28), suggesting that at least some of the sites of BG4 CUT&Tag enrichment could be the result of Tn5 cleavage at accessible chromatin. The strong genome-wide correlation between ATAC-seq, G4Access, BG4 CUT&Tag, and untargeted CUT&Tag (**Figure 5C**) suggests that chromatin accessibility strongly dictates enrichment of both BG4-targeted and untargeted CUT&Tag signal. This is also not surprising, as G4s form in ssDNA within nucleosome-depleted chromatin, and G4 formation may actively exclude nearby nucleosomes (5, 60, 61). Third, other literature has implicated both hyperactive Tn5 (62) and Protein A (63) in binding to G4s, suggesting that untargeted cleavage may also occur due to unanticipated G4 binding by transposase components at existing G4s.

When taken as a whole, our findings suggest that stringent biochemical validation of sites of BG4 CUT&Tag enrichment should always be performed to strengthen the confidence in BG4 CUT&Tag analyses, such as knockdown of G4-resolving enzymes followed by an examination of BG4 CUT&Tag signal increase (as in (22)), confirmation of individual G4s with circular dichroism spectroscopy (reviewed in (64)), and/or *in vitro* binding assays examining direct interaction between the examined G4s and other colocalizing proteins. However, as validation of every peak of enrichment across an entire genome is not realistic to perform, this lessens our confidence in BG4 CUT&Tag analyses in the absence of other corroborating data.

## DATA AVAILABILITY

All data utilized for this analysis is available from GEO (10), SRA (11), or ENCODE (29) at the accessions included in **Supplementary Tables 1, 2, and 3**.

## SUPPLEMENTARY DATA

Supplementary Data are available at NAR online.

## AUTHOR CONTRIBUTIONS

MDT: Analysis, Methodology, Writing-original draft. AKB: Conceptualization, Writing—review & editing.

