## Supplementary Information for "Untargeted CUT&Tag and BG4 CUT&Tag are both enriched at G-quadruplexes and accessible chromatin"

**Supplementary Table 1.** Untargeted CUT&Tag datasets from GEO (1).

| Cell Type | Antibody | GEO Accession | SRA Accession | Source Publication |
|---|---|---|---|---|
| MOLM13 | Non-targeting IgG | GSM7101045 | SRR23876011 | (2) |
| HepG2 | Non-targeting IgG | GSM6306788 | SRR20020486 | (3) |
| Karpas422 | Non-targeting IgG | GSM5558304 | SRR15729258 | (4) |
| Farage | Non-targeting IgG | GSM5558296 | SRR15729250 | (4) |
| HCT-116 | Non-targeting IgG | GSM6235290 | SRR19631169 | (5) |
| Human umbilical cord mesenchymal stem cells | Non-targeting IgG | GSM7070802 | SRR23641484 | (6) |
| PC-3 | Non-targeting IgG | GSM6919287 | SRR22990288 | (7) |
| Lung lymphatic endothelial cells | Non-targeting IgG | GSM6806707 | SRR22577939 | (8) |
| J-Lat 10.6 | Non-targeting IgG | GSM6590675 | SRR21625457 | (9) |
| HEK293T | Non-targeting IgG | GSM6504650 | SRR21211482 | (10) |
| Kasumi-1 | Non-targeting IgG | GSM5504185 | SRR15353246 | (11) |
| THP-1 | Non-targeting IgG | GSM5504188 | SRR15353249 | (11) |
| Cal27 | Non-targeting IgG | GSM5641128 | SRR16502256 | (12) |
| FHs74Int | Non-targeting IgG | GSM6574214 | SRR21661660 | (13) |
| HEK293-T | Non-targeting IgG | GSM6267563 | SRR19859937 | (14) |
| H1 | Non-targeting IgG | GSM3560256 | SRR8435038 | (15) |
| K562 | Non-targeting IgG | GSM3560264 | SRR8435051 | (15) |
| U2OS | None | GSM5501191 | SRR15337933 | (16) |
| K562 | None | GSM5501188 | SRR15337930 | (16) |
| MCF7 | None | GSM5501227 | SRR15337969 | (16) |
| H1 | None | GSM6627988 | SRR21859393 | (17) |

**Supplementary Table 2.** Other GEO (1) Datasets used in this study. Biological replicates are indicated, with technical replicates indicated when necessary in parentheses.

| Method | Target | Cell Type | Replicate | GEO Accession | SRA Accession | File Type | Source Publication |
|---|---|---|---|---|---|---|---|
| CUT&Tag | BG4 | K562 | 1 (T1) | GSM5501186 | N/A | .bw | (16) |
| CUT&Tag | BG4 | K562 | 1 (T2) | GSM5501187 | N/A | .bw | (16) |
| CUT&Tag | BG4 | K562 | 2 (T1) | GSM5501192 | N/A | .bw | (16) |
| CUT&Tag | BG4 | K562 | 2 (T2) | GSM5501193 | N/A | .bw | (16) |
| CUT&Tag | BG4 | K562 | 3 (T1) | GSM5501198 | N/A | .bw | (16) |
| CUT&Tag | BG4 | K562 | 3 (T2) | GSM5501199 | N/A | .bw | (16) |
| CUT&Tag | BG4 | K562 | 1, 2, 3 | GSE181373 | N/A | .bed | (16) |
| CUT&Tag | Non-targeting IgG | K562 | 1 | GSM5501188 | N/A | .bw | (16) |
| CUT&Tag | Non-targeting IgG | K562 | 2 | GSM5501194 | N/A | .bw | (16) |
| CUT&Tag | Non-targeting IgG | K562 | 3 | GSM5501200 | N/A | .bw | (16) |
| ChIP-seq | BG4 | K562 | 1 | GSM4948705 | SRR13161747 | .fastq | (18) |
| ChIP-seq | BG4 | K562 | 1, 2 | GSE162299 | N/A | .bed | (18) |
| ChIP-seq | Input | K562 | 1 | GSM4948764 | SRR13161615 | .fastq | (18) |
| G4Access | G4 | K562 | 1 (T1) | GSM5665726 | SRR16700952 | .fastq | (19) |
| G4Access | G4 | K562 | 1 (T2) | GSM5665727 | SRR16700953 | .fastq | (19) |
| G4Access | G4 | K562 | 1 (T3) | GSM5665728 | SRR16700954 | .fastq | (19) |
| G4Access | G4 | K562 | 2 | GSM5665729 | SRR16700955 | .fastq | (19) |
| G4Access | Input | K562 | 1 | GSM5665730 | SRR16700956 | .fastq | (19) |
| CUT&Tag | H3K4me3 | K562 | 1 | GSM3536518 | SRR8383516 | .fastq | (15) |
| CUT&Tag | H3K27me3 | K562 | 1 | GSM3560261 | SRR8435047 | .fastq | (15) |
| CUT&Tag | Non-targeting IgG | K562 | 1 | GSM3560264 | SRR8435051 | .fastq | (15) |
| CUT&Tag | BG4 | U2OS | 1 (T1) | GSM5501189 | N/A | .bw | (16) |
| CUT&Tag | BG4 | U2OS | 1 (T2) | GSM5501190 | N/A | .bw | (16) |
| CUT&Tag | BG4 | U2OS | 2 (T1) | GSM5501195 | N/A | .bw | (16) |
| CUT&Tag | BG4 | U2OS | 2 (T2) | GSM5501196 | N/A | .bw | (16) |
| CUT&Tag | BG4 | U2OS | 3 (T1) | GSM5501201 | N/A | .bw | (16) |
| CUT&Tag | BG4 | U2OS | 3 (T2) | GSM5501202 | N/A | .bw | (16) |
| CUT&Tag | BG4 | U2OS | 1, 2, 3 | GSE181373 | N/A | .bed | (16) |
| CUT&Tag | No IgG | U2OS | 1 | GSM5501191 | N/A | .bw | (16) |
| CUT&Tag | No IgG | U2OS | 2 | GSM5501197 | N/A | .bw | (16) |
| CUT&Tag | No IgG | U2OS | 3 | GSM5501203 | N/A | .bw | (16) |
| ChIP-seq | BG4 | U2OS | 1 | GSM5222763 | SRR14129547 | .fastq | (18) |
| ChIP-seq | BG4 | U2OS | 1, 2 | GSE162299 | N/A | .bed | (18) |
| ChIP-seq | Input | U2OS | 1 | GSM5222766 | SRR14129553 | .fastq | (18) |
| ATAC-seq | N/A | U2OS | 1 (T1) | GSM3462884 | SRR8171329 | .fastq | (20) |
| ATAC-seq | N/A | U2OS | 1 (T2) | GSM3462883 | SRR8171328 | .fastq | (20) |
| ATAC-seq | N/A | U2OS | 1 (T3) | GSM3462882 | SRR8171327 | .fastq | (20) |
| ATAC-seq | N/A | U2OS | 1 (T4) | GSM3462881 | SRR8171326 | .fastq | (20) |
| ATAC-seq | N/A | U2OS | 2 (T1) | GSM3462880 | SRR8171325 | .fastq | (20) |
| ATAC-seq | N/A | U2OS | 2 (T3) | GSM3462878 | SRR8171323 | .fastq | (20) |
| ATAC-seq | N/A | U2OS | 2 (T4) | GSM3462877 | SRR8171322 | .fastq | (20) |
| ATAC | N/A | U2OS | 1, 2 | GSE121840 | N/A | .bed | (20) |

**Supplementary Table 3**. ENCODE (21) datasets used in this study.

| Method | Target | Cell Type | Biological Replicate | Accession (ENCODE) | File Type |
|---|---|---|---|---|---|
| ATAC-seq | N/A | K562 | 1 | ENCSR868FGK | .bam |
| ATAC-seq | N/A | K562 | 2 | ENCSR868FGK | .bam |
| ATAC-seq | N/A | K562 | 3 | ENCSR868FGK | .bam |
| ATAC-seq | N/A | K562 | 1, 2, 3 | ENCFF057UYP | .bed |
| ChIP-seq | H3K4me3 | K562 | 1, 2 | ENCFF403DTU | .bed |
| ChIP-seq | H3K27me3 | K562 | 1, 2 | ENCFF795ZOS | .bed |

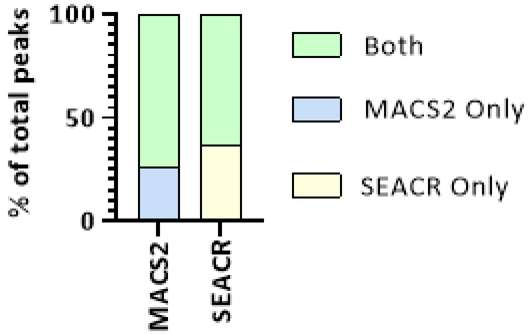


**Supplementary Figure 1.** The majority of untargeted CUT&Tag peaks are identified by both MACS2 and SEACR. Percentage of overlap of untargeted CUT&Tag peaks shared called by MACS2 (22) (n = 4221) and SEACR (23) (n = 4075).

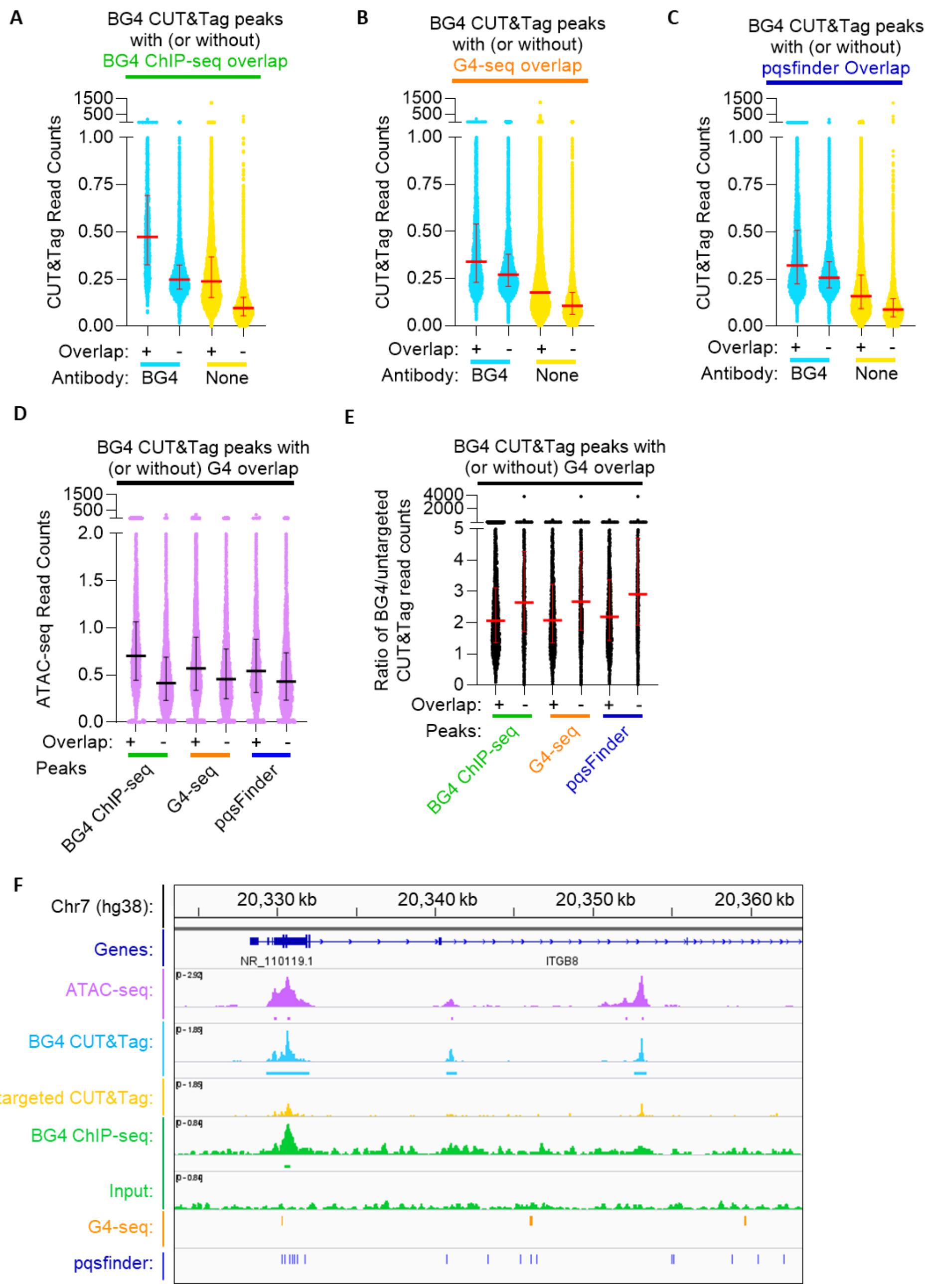


**Supplementary Figure 2.** Enrichment of BG4 CUT&Tag over untargeted CUT&Tag is not increased at G4s in U2OS cells. (**A-D**). BG4 or untargeted (no IgG) CUT&Tag signal read counts were quantified at CUT&Tag peaks that overlap or do not overlap a G4 mapped by BG4 ChIP-seq (18) (**A**), G4-seq (24) (**B**), or pqsfinder (25) (**C**) in U2OS cells. Data are the average of 3 biological replicates, and the median and interquartile range is plotted in red. (**D**) ATAC-seq (20) read counts quantified at BG4 CUT&Tag peaks that overlap or do not overlap a G4 mapped by the indicated method. Data are the average of 2 biological replicates, and the median and interquartile range is plotted in black. (**E**) Ratio of read counts of BG4 (16) and untargeted CUT&Tag from U2OS cells (16) quantified at CUT&Tag peaks which overlapped or did not overlap a G4 mapped by the indicated method. Data are the average of 3 biological replicates, and the median and interquartile range is plotted in red. (**F**) Read counts of ATAC-seq (20), BG4 CUT&Tag (16), untargeted CUT&Tag (16), BG4 ChIP-seq (18), and ChIP-seq input (18) from U2OS cells. Also shown are G4-seq-mapped G4s (24) on the indicated strand and pqsfinder-predicted G4s (25) at representative locus on chromosome 1. ATAC-seq read counts are the average of 2 biological replicates. BG4 CUT&Tag and untargeted CUT&Tag read counts are the average of 3 biological replicates. BG4 ChIP-seq and ChIP-seq input are 1 biological replicate.

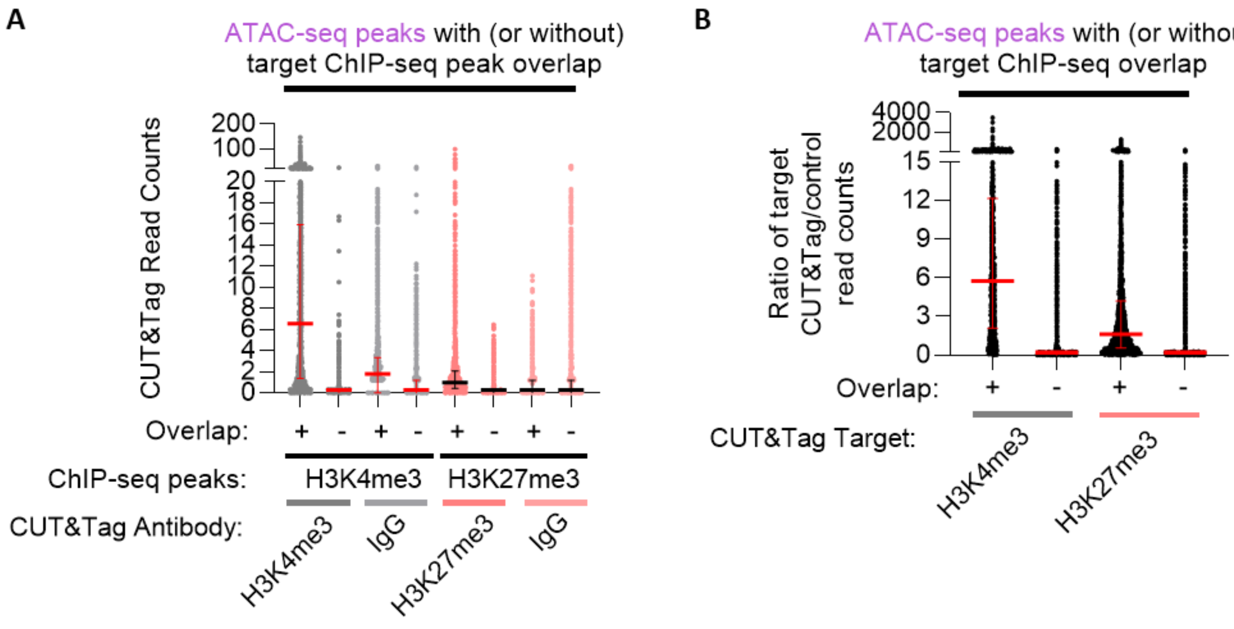


**Supplementary Figure 3.** Enrichment of histone PTM CUT&Tag over untargeted CUT&Tag within accessible chromatin increases in the presence of a ChIP-seq mapped histone PTM site. (**A**) H3K4me3, H3K27me3, and negative control (non-targeting IgG) CUT&Tag read counts from K562 cells (15) were quantified at ATAC-seq peaks that overlap or do not overlap H3K4me3 (ENCSR668LDD) or H3K27me3 (ENCSR000EWB) ChIP-seq peaks. Read counts are for 1 biological replicate; the median and interquartile range is plotted in red and black for H3K4me3 and H3K27me3, respectively. (**B**) Ratio of read counts of H3K4me3 or H3K27me3 CUT&Tag over untargeted CUT&Tag from K562 cells (15) at ATAC-seq peaks that overlap or do not overlap H3K4me3 (ENCSR668LDD) or H3K27me3 (ENCSR000EWB) ChIP-seq peaks. Read counts are for 1 biological replicate; the median and interquartile range is plotted in red.

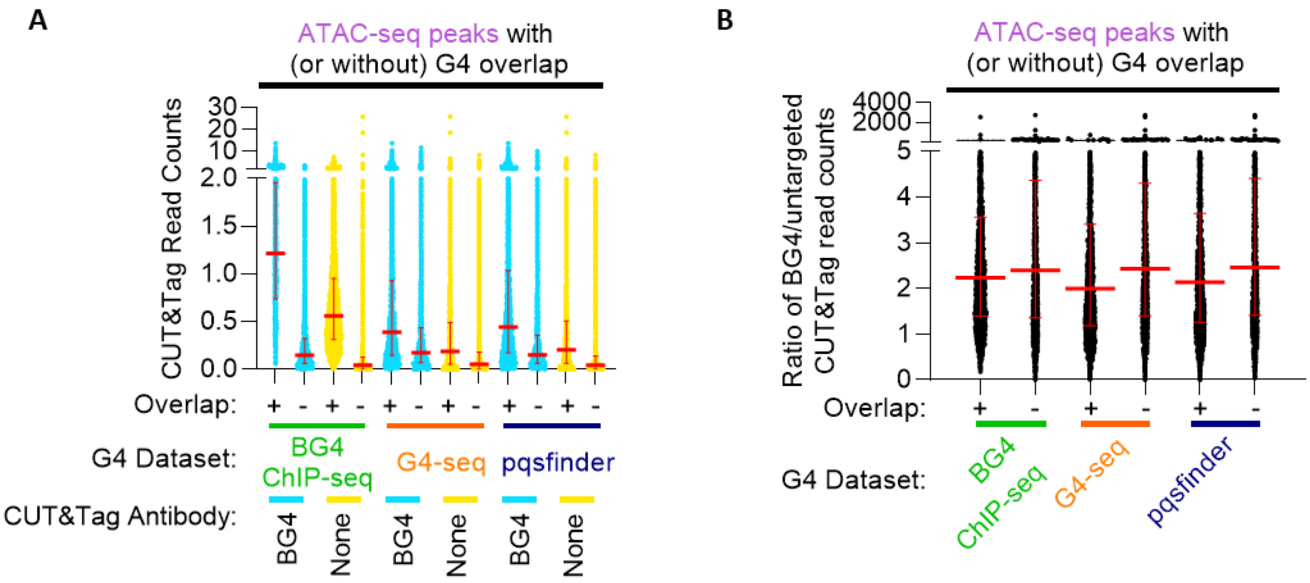


**Supplementary Figure 4.** The ratio of read counts from BG4 CUT&Tag over untargeted CUT&Tag at individual peaks within accessible chromatin does not rise in the presence of a mapped G4 in U2OS cells. (**A**) Read counts of BG4 CUT&Tag (16) and untargeted CUT&Tag (16) from U2OS cells at ATAC-seq peaks (20) that overlap or do not overlap a mapped G4 (18, 24, 25). Data are the average of 3 biological replicates. The median and interquartile range is plotted in red (**B**) Ratio of read counts of BG4 CUT&Tag (16) and untargeted CUT&Tag (16) from U2OS cells quantified at ATAC-seq peaks (20) that overlap or do not overlap a G4 mapped by the indicated method (18, 19, 24, 25). Data are the average of 2 biological replicates. The median and interquartile range is plotted in red.
